# Investigating Predictive Coding in Younger and Older Children Using MEG and a Multi-Feature Auditory Oddball Paradigm

**DOI:** 10.1101/2022.07.26.501632

**Authors:** Hannah Rapaport, Robert A. Seymour, Nicholas Benikos, Wei He, Elizabeth Pellicano, Paul F. Sowman

## Abstract

There is mounting evidence for predictive coding theory from computational, neuroimaging, and psychological research. However there remains a lack of research exploring how predictive brain function develops across childhood. To address this gap, we used paediatric magnetoencephalography (MEG) to record the evoked magnetic fields of 18 younger children (*M* = 4.1 years) and 19 older children (*M* = 6.2 years) as they listened to a 12-minute auditory oddball paradigm. For each child, we computed a mismatch field ‘MMF’: an electrophysiological component that is widely interpreted as a neural signature of predictive coding. Consistent with our hypotheses, the older children showed significantly larger MMF amplitudes relative to the younger children. Furthermore, the older children showed a significantly larger MMF amplitude in the right inferior frontal gyrus (IFG; 0.312 to 0.33 s) relative to the younger children, *p* < .05. These findings support the idea that predictive brain function develops during childhood, with increasing involvement of the frontal cortex in response to prediction errors. These findings contribute to a deeper understanding of the brain function underpinning child cognitive development.

**Highlights:**

- This is the first paediatric MEG study to examine the sources underlying the MMF.
- Older children showed larger MMF amplitudes in the right inferior frontal gyrus.
- Results support the idea that predictive brain function develops during childhood.

## 1. Introduction

Under predictive coding theory, the brain houses an internal, probabilistic, generative model which represents the statistical structure of the external world (Clark, 2015; Friston & Kiebel, 2009; Hohwy, 2013; Rao & Ballard, 1999). The brain uses this model to generate predictions about the most likely causes of incoming sensory signals, and tests its model by comparing its top-down prior predictions against bottom-up sensory signals from the world (Hohwy, 2012). Predictions and sensory signals can be represented as probability distributions whereby the difference between the two represents the model’s ‘prediction error’. Prediction error is a useful learning signal as it indicates which sensory information the model failed to predict and hence, which information requires further processing at higher levels of the neural hierarchy (Dołęga & Dewhurst, 2021). To reduce processing requirements, the brain strives to minimise prediction errors over time (Clark, 2013).

An account of how children come to know about the world falls naturally from predictive coding principles (Badcock et al., 2019; Gopnik, 2012; Scholl, 2005). Infants are thought to begin life with a set of innately specified priors, rooted in the genes. Following environmental stimulation, ‘innate priors’ are updated through ‘perceptual inference’ (Hohwy, 2012) and subsequently called ‘empirical priors’. Thus, predictive coding takes advantage of both nature and nurture perspectives, allowing them to interact within a unifying framework. Iterative model updating should enable children to meet incoming sensory signals with increasingly accurate priors which should, in turn, result in fewer prediction errors, and hence greater confidence in the model’s predictions. Children’s priors may become increasingly precise as they gain experience in the world (Kayhan et al., 2019; Köster et al., 2020). As such, the posterior distribution (i.e., the perceptual experience; Hohwy, 2012) would be gradually biased towards the increasingly narrow prior distribution (representing the brain’s stored knowledge), and away from the sensory signal distribution (Lucas et al., 2014).

Rapid model maturation across childhood may be underpinned by concurrent and significant neurophysiological changes. Indeed, the brain roughly quadruples in weight before age six, by which time it has reached approximately 90% of its adult volume (Brown & Jernigan, 2012). Furthermore, the efficiency of neuronal communication increases due to a prolonged period of synaptic pruning and myelination during childhood and adolescence (Blakemore & Choudhury, 2006; Fischer & Bidell, 2007; Santos & Noggle, 2011). In particular, the protracted maturation of the prefrontal cortex (Huttenlocher & Dabholkar, 1997)—a region thought to play a key role in extracting complex statistical regularities from incoming sensory signals (Basirat et al., 2014; Dürschmid et al., 2016) — may support increasingly sophisticated predictive brain function across development (Krogh et al., 2013). While a predictive coding account of neurocognitive development is supported by an extensive body of behavioural evidence (Gopnik & Wellman, 2012; Köster et al., 2020), neural evidence is lacking (Emberson et al., 2019; Zhang & Emberson, 2020)—partly due to practical challenges associated with conducting function brain recordings with young children (Barkovich et al., 2019). We sought to address this gap in the literature.

A popular method for testing predictive coding is to use electroencephalography (EEG) and/or magnetoencephalography (MEG) to record participants’ brain responses as they listen to an auditory oddball paradigm (Friston, 2005; Heilbron & Chait, 2017). These paradigms are comprised of high-probability ‘standard’ and lower-probability ‘deviant’ stimuli (e.g., pure tones). Averaged evoked responses to the ‘standards’ and ‘deviants’ typically diverge between 0.1 and 0.25 s following stimulus onset, with the ‘deviant’ showing a larger amplitude relative to the ‘standard’ waveform (Näätänen et al., 2019b). This divergence—conventionally presented as a difference waveform—is the ‘mismatch negativity’ (MMN; EEG literature) or the ‘mismatch field’ (MMF; MEG literature; hereafter, the MMN/F).

The MMN/F has been interpreted as a neural index of prediction error (Friston, 2005), whereby the larger the evoked response amplitude, the larger the corresponding prediction error signal. With significant neurophysiological maturation during childhood (Blakemore & Choudhury, 2006; Brown & Jernigan, 2012; Huttenlocher & Dabholkar, 1997), children’s brains may become increasingly proficient at extracting statistical regularities from the auditory input (Saffran et al., 2001). Consequently, older children may be able to form relatively precise priors of the upcoming stimuli, whereby the strongest prediction would be for the presentation of a high-probability ‘standard’. Thus, the presentation of a lower-probability ‘deviant’ may evoke a larger prediction error relative to that of a ‘standard’. This, in turn, would give rise to a relatively large MMF amplitude. By contrast, if younger participants are less proficient at extracting these statistical regularities, then they may form comparatively less-precise predictions for the upcoming stimuli. Thus, if younger children are less-precisely predicting the presentation of a standard, then the presentation of both high-probability ‘standards’ and lower-probability ‘deviants’ may evoke similar degrees of prediction error, resulting in a relatively attenuated MMF.

Overall, one might expect a larger MMF amplitude in older relative to younger children, reflecting maturation of predictive brain function across development. However, previous findings have been mixed, with studies reporting an increase (Bishop, Anderson, et al., 2011; Chobert et al., 2014; Linnavalli et al., 2018; Oades et al., 1997; Putkinen, Tervaniemi, Saarikivi, de Vent, et al., 2014; Putkinen, Tervaniemi, Saarikivi, Ojala, et al., 2014), decrease (Csépe et al., 1992; Kraus, McGee, Carrell, et al., 1993; Kraus, McGee, Micco, et al., 1993), or no difference (Gomot et al., 2000; Kraus et al., 1999; Shafer et al., 2000, 2010) in the MMN amplitude with age.

Here we aimed to test a predictive coding account of early neurocognitive development. To this end, we used paediatric MEG (Johnson et al., 2010; Rapaport et al., 2019) and a multi-feature auditory oddball paradigm, and measured MMF responses in younger (*M*_age_ = 4.1 years) and older (*M*_age_ = 6.2 years) children. We hypothesised that the older-relative-to-younger children would show a larger MMF amplitude, reflecting maturation of predictive brain function across development. Furthermore, we hypothesised that this maturation would be underpinned by increasing involvement of the frontal cortex in responding to prediction errors, as reflected by a significantly larger frontal-MMF in the older children.

## 2. Methods

### 2.1. Participants

Sixty children (aged 3 to 6 years) were recruited via the Macquarie University ‘Neuronauts’ child research participation database. Of those, three children were unable to complete the MEG recording session due to anxiety (*n* = 2, aged 3.1 and 5.5 years) or falling asleep during the recording (*n* = 1, aged 3.1 years). Of the 57 collected datasets, 20 (35%) were excluded due to: (a) having a poor head position in relation to the MEG sensor array (*n* = 6; determined based on visual inspection), (b) technical issues (e.g., malfunctioning MEG channels, event marking, or real-time head tracking coils; *n* = 10), and (c) excessive in-scanner head motion (i.e., exceeding an average of 10 mm of movement over the course of the recording; *n* = 4; *M*_age_ = 4.7 years, range = 3.2–5.8). Thus, the final sample included 37 children.

To investigate the (cross-sectional) developmental trajectory of the MMF, we performed a median age split—separating the data groups of younger children (*n* = 18; *M*_age_ = 4.1 years, SD = 0.9, range = 3.1–5.3 years; 8 females; 3 ambidextrous, 3 left-handed) and older children (*n* = 19; *M*_age_ = 6.2 years, SD = 0.4, range = 5.6–6.9; 12 females; 2 ambidextrous, 2 left-handed determined based on parent report). Normal hearing thresholds between 500 Hz and 1500 Hz were confirmed with pure-tone audiometric testing using an Otovation Amplitude T3 series audiometer (Otovation LLC, PA, United States). All children had parent-reported normal or corrected-to-normal vision and had no history of developmental disorders, epilepsy, brain injury or language or speech impairment, as reported by parents. Children and their parents provided verbal and written informed consent, respectively, before the experiment. All procedures were approved by the Macquarie University Human Research Ethics Committee (reference: 5201600188). Families were paid 40 AUD for their participation and the children received a gift bag and certificate.

### 2.2. Auditory Oddball Paradigm

Electrophysiological responses were measured as participants listened to a passive multi-feature auditory oddball paradigm (adapted from Näätänen et al., 2004). We chose to use the multi-feature over the traditional oddball paradigm as the former version offers advantages in terms of time efficiency—a quality which is of critical importance when testing young children who are generally less compliant than adults (Lovio et al., 2009). The multi-feature recording time is significantly reduced by presenting standards and deviants in an alternating pattern, as opposed to the traditional paradigm whereby the ratio of deviants to standards is between 1:7 to 1:10 (Petermann et al., 2009). No differences have been found between MMNs elicited by multi-feature compared to traditional oddball paradigms (Näätänen et al., 2004; Pakarinen et al., 2007). Furthermore, the multi-feature paradigm has been shown to reliably evoke mismatch responses in children (Petermann et al., 2009; Putkinen et al., 2012; Putkinen, Tervaniemi, Saarikivi, Ojala, et al., 2014).

The paradigm consisted of three sequential stimulus blocks, each of which ran for approximately four minutes and consisted of 495 stimuli (0.5 s stimulus-onset-asynchrony). Block order was counterbalanced across participants. Each block began with the presentation of 15 ‘standard’ stimuli followed by alternating ‘standard’ and ‘deviant’ stimuli (see Figure 1 for a schematic diagram). The ‘standard’ stimuli were sinusoidal harmonic tones which were 75 ms in duration (including 5 ms rise and fall times) and presented at a volume of 80 SPL and at a frequency of 550, 1000 and 1500 Hz (for each block, respectively). Each ‘standard’ had a presentation probability of P = 0.5 (excluding the first 15 standards in each block).

**Figure 1.**
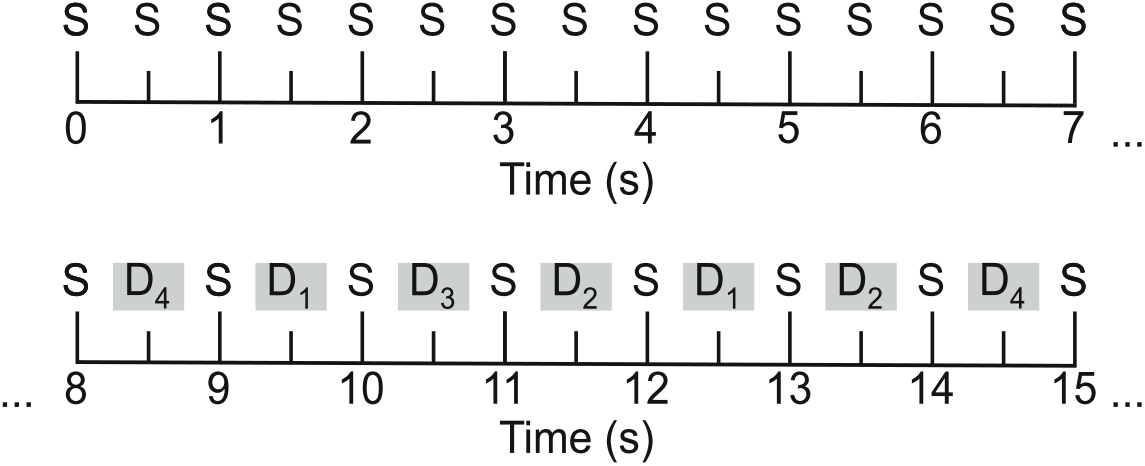
A schematic illustration of the multi-feature auditory oddball paradigm. The first 15 tones are ‘standards’ (S) followed by alternating ‘standard’ and one of four types of ‘deviant’ (D) tones.

The ‘deviant’ stimuli differed from the ‘standards’ in one of four acoustic features: (1) frequency (half were 10% higher [partials: 605, 1100, 1650 Hz] and half were 10% lower [partials: 495, 900, 1350 Hz] than the standards), (2) intensity (half were 10% louder and half were 10% softer than the standards), (3) duration (the stimulus was presented for 25 ms with a silent 25 ms gap before the next stimulus), or (4) by having a silent gap in the middle of the stimulus (i.e., 7 ms removed from the middle of the stimulus tone, including 1 ms fall and rise times). Each of the four deviant subtypes had a presentation probability of P = 0.125. Deviants were presented in a pseudorandom order such that the same type of deviant could not be presented in succession. The paradigm was programmed and presented in MATLAB (Mathworks, Natick, MA, USA) using Psychtoolbox (Brainard, 1997).

### 2.3. MEG Acquisition

Electrophysiological data were acquired using a whole-head, supine, paediatric MEG system (Model PQ1064R-N2m, KIT, Kanazawa, Japan) housed in a magnetically shielded room (MSR; Fujihara Co. Ltd., Tokyo, Japan). The MEG sensor array contained 125 first-order axial gradiometers, each of which had a coil diameter of 15.5 mm and a baseline of 50 mm (see He et al., 2019 for further details). The dewar was designed to fit a maximum head circumference of 53.4 cm, accommodating the heads of more than 90% of 5-year-old Caucasian children (Johnson et al., 2010). All data were acquired at a sampling rate of 1000 Hz and with an online bandpass filter of 0.03–200 Hz. To maximise the likelihood of obtaining high-quality data, we followed a child-friendly MEG testing protocol (see Rapaport et al., 2019).

Before the MEG recording, participants were fitted with a polyester cap containing five head position indicator (HPI) ‘marker’ coils. A digitiser pen (Polhemus Fastrak, Colchester, USA) was used to record the locations of the HPI coils, as well as three fiducial points (the nasion and bilateral pre-auricular points) and 300-500 points from the scalp and face. During the MEG recording, participants listened to the multi-feature paradigm whilst watching a silent video of their choice. The paradigm was presented via a 60 cm^2^ speaker (Panphonics SSH sound shower, Panphonics) positioned centrally at the foot of the MEG bed. The video was projected onto the MSR ceiling above the dewar. Participants’ head position in relation to the MEG sensor array was continuously monitored using a real-time marker coil tracking system (Oyama et al., 2012). Participants were accompanied by a researcher (and often a caregiver) during the MEG recording. The MEG procedures, including the set-up and the recording, took approximately 45 minutes to complete.

### 2.4. Data Pre-processing

Following an inspection of the data, we removed three MEG channels that were consistently flat and noisy across participants for more than 10% of the recording (Gross et al., 2013). The following steps were performed MEG160 (Yokogawa Electric Corporation and Eagle Technology Corporation, Tokyo, Japan). Environmental noise—estimated based on recordings from three reference magnetometers—was suppressed using a Time-Shift Principal Component Analysis (TSPCA) algorithm (de Cheveigné & Simon, 2007; block width: 10,000 ms, 3 shifts). Data acquired with the real-time marker coil tracking system were used to correct for head motion artefacts (Knösche, 2002; realignment conditions: sphere mesh = 321, prune ratio = 0.05). Head motion artefact correction could not be performed for seven datasets that had missing or erroneous real-time coil tracking data for more than 10% of the recording. These seven datasets were still included as head movement between the pre- and post-recording marker coil measurements did not exceed 5 mm.

Further pre-processing steps were performed in Matlab 2020a (MathWorths, Inc., Natick, MA, USA) using the Fieldtrip Toolbox (v20200213; Oostenveld et al., 2011). For each participant, the entire recording was high- and low-pass filtered at 0.1 and 40 Hz, respectively (using a onepass-zerophase firws filter with a Blackman window), and band-stop filtered to remove residual 50 Hz power-line contamination and its harmonics. Following visual inspection, segments of the recording containing artefacts (e.g., SQUID jumps and jaw clenches) were removed. Channels that contained a large number of these visually-identified artefacts and/or were flat for more than 10% of the recording were interpolated to ensure that all participants had the same number of channels (Medvedovsky et al., 2007; NB. these removals were in addition to the three channels already rejected across the entire sample). An independent component analysis (ICA) was used to suppress eye blink artefacts: the raw recordings were high-pass filtered at 1Hz to improve the ICA performance (Winkler et al., 2015) and then components with scalp distributions that corresponded to eye blinks were removed from the 0.1 Hz, pre-processed data. Up to one independent component was removed per participant.

The continuous data were epoched into segments of 0.5 s (0.1 s pre-, and 0.4 s post-stimulus onset). The first 15 epochs of each stimulus block were excluded from further analysis. Standard and deviant epochs were averaged across, respectively, to compute standard and deviant event-related fields (ERFs). It should be noted that we averaged across all four deviant conditions, as: (1) we sought to maximise the statistical power of our comparisons, and (2) we had no theoretical reason to analyse the four types of deviants separately. While the deviants differed in terms of their acoustic features, the constant feature that unites them is their pseudorandom occurrence (in contrast to the relatively predictable occurrence of the standards – see Figure 1). The standard ERF was subtracted from the deviant ERF to produce a mismatch field (MMF).

### 2.5. Sensor-Level Analysis

The sensor-level analysis involved two key steps performed in MNE-Python (Gramfort et al., 2013): (1) identifying a time-window of interest (TOI), and (2) constraining the age-related analysis to this time-window. We used a data-driven approach to identify a TOI. First, we reduced the multi-variate (channel x time) data from all 37 participants into a single time-course—achieved using an ‘Effect-Matched Spatial (EMS) filtering’ approach (Schurger et al., 2013). EMS filtering involves estimating a spatial filter from the data itself, and then projecting the data through the spatial filter. EMS is superior to other data reduction methods (e.g., averaging a contiguous cluster of channels) as it accounts for the spatiotemporal evolution of the experimental effect across time and space, and weights the channels at each timepoint accordingly. To avoid circularity in the procedure, we employed five-fold stratified cross-validation procedure (Engemann & King, 2021; Huang et al., 2020). This analysis involved repeating the following steps four times for each participant:

1. The data were normalised using z-scores.
2. A spatial filter was derived from 80% of the dataset, leaving out a different 20% for each repetition. The spatial filter was computed by subtracting the ‘standard’ from the ‘deviant’ epochs at each timepoint and channel. This produced a set of weightings (i.e., a spatial filter) that reflected the magnitude of the experimental effect (i.e., the MMF) over temporal and spatial dimensions.
3. The spatial filter was applied to the remaining 20% of the dataset by taking the product of the filter and the data at each time point and for each channel, and summing the results to render a single, spatially filtered time-course for the standard and deviant conditions, respectively.

The 37 time-courses were then averaged to form a single time-course for the ‘standard’ and ‘deviant’ conditions, respectively. Subsequently, we performed a non-parametric cluster-based permutation analysis (Maris & Oostenveld, 2007) on the whole-group EMS filtered data to determine the time-course of the MMF effect. This permutation approach has been shown to adequately control the Type-I error rate for electrophysiological data. The analysis involved conducting one-sample *t*-tests at each time point to determine whether the MMF amplitude was significantly different from zero. We clustered samples whose *t*-values fell below a threshold corresponding to an alpha level of 0.05 (on the basis of temporal proximity) and calculated cluster-level test statistics by taking the average of the *t*-values within each cluster. The data were then permuted 1,000 times, each time randomly shuffling the ‘standard’ and ‘deviant’ condition labels and recomputing the *t*-values. We constructed a permutation distribution from these random partition *t*-values. Finally, the significance of each cluster determined by using a threshold Monte-Carlo *p*-value.

To maximise statistical power, the time window of the significant cluster (identified in the analysis above) was used to constrain the age-related analysis. First, for each participant, we computed a mean MMF^1^ amplitude by taking the average of their EMS-filtered data within the TOI.

Here the MMF was liberally defined as any significantly larger deviant-relative-to-standard amplitude within the time window of interest. Our definition stands in contrast to the definition of the classic adult MMN/F, which emerges soon after the second ‘N1’ or ‘M2’ component (Hari & Puce, 2017, p. 265; Näätänen et al., 2019a, p. 53) and is visible between 0.1 to 0.25 s following stimulus onset (Näätänen et al., 2019b). We relied on this more liberal definition as the latency of the peak mismatch effect changes across childhood (Näätänen et al., 2019a) and it would therefore have been inappropriate to constrain our analysis to the classic adult MMN time window. It should be noted that prior paediatric auditory oddball studies have likewise used liberally-defined time windows to extract MMN effects (e.g., 0.2– 0.33 s in 6- to 7-year-olds, Lovio et al., 2009; 0.3–0.5 s in 4- to 12-year-olds, Partanen et al., 2013; 0.15–0.4 s in 5- to 7-year-olds, Petermann et al., 2009; 0.1–0.3 s in 9- to 13-year-olds, Putkinen et al., 2014; 0.1–0.32 s in 4- to 10-year-olds, Shafer et al., 2000).

We compared mean MMF amplitudes between the younger and older children using a two-sided permutation independent-samples *t*-test with 5000 bootstrap samples. For each *p*-value, we performed 5,000 reshuffles of the age group labels using the DABEST (Data Analysis with Bootstrap-coupled ESTimation; Ho et al., 2019) open-source libraries for Python.

### 2.6. Source-Level Analysis

To investigate how the neural network underpinning the MMF changes across age, we conducted source analysis on six predefined regions of interest (ROIs): bilateral primary auditory cortices (A1), bilateral superior temporal gyri (STG), and bilateral inferior frontal gyri (IFG; Garrido et al., 2007, 2008, 2009; Phillips et al., 2015). These ROIs were drawn from the adult literature as this is, to our knowledge, the first paediatric MEG study which has examined the neural sources underpinning MMN/F generation in typically developing children.

As the participants did not have individual structural MRI scans, we used the MRI Estimation for MEG Sourcespace toolbox (MEMES; Seymour, 2018; https://github.com/Macquarie-MEG-Research/MEMES) to create surrogate structural MRIs. This procedure uses an Iterative Closest Point algorithm to match the participant’s digitised head shape information to an age-appropriate average MRI template (Neurodevelopmental MRI Database; Richards et al., 2016).

For each participant, the best-fitting template with the lowest objective registration (Gohel et al., 2017) was selected and used to create a cortical mesh and source grid and co-registered with the MEG sensor locations. A forward model (i.e., leadfield) was then computed using the cortical mesh as the volume conductor model. Source reconstruction was performed using a linearly constrained minimum variance (LCMV) beamformer (Van Veen et al., 1997), as implemented in Fieldtrip Toolbox (v20200213; Oostenveld et al., 2011). A spatial filter for each vertex in the source grid was computed using a free dipole orientation (Garrido et al., 2007). This step was performed separately for the ‘standard’ and ‘deviant’ conditions based on the covariance matrix calculated from the data combined across conditions. The spatial filters for all vertices within each ROI were combined into a single spatial filter by weighting voxels within the ROI according to their proximity to a centroid (Brookes et al., 2016). This centroid was defined as the voxel within the ROI that was nearest to all other voxels in the ROI (Douw et al., 2018). The sensor-level data was then right-multiplied by the spatial filters for each condition.

The final two analysis steps involved using the non-parametric cluster-based permutation approach. For both tests, we permuted the data 2,000 times (Maris & Oostenveld, 2007). First, we used paired-samples *t-*tests to compare the ‘standard’ and ‘deviant’ amplitudes at each of the ROIs for all 37 children. Second, we used independent-samples *t*-tests to compare the source-level activity between the groups of younger and older children. We permuted the condition labels (‘standard’ and ‘deviant’) for the paired tests, and group labels (‘younger’ and ‘older) for the independent tests.

The experimental paradigm, data pre-processing and data analysis scripts are freely available on an Open Science Framework repository: https://osf.io/35q8n/.

## 3. Results

### 3.1. Sensor-Level Results

Across all 37 children, we found a significantly larger deviant-relative-to-standard (i.e., MMF) amplitude between 0.19 and 0.49 s following stimulus onset (see Figure 2). Results from a two-sided permutation independent-samples *t*-test (constrained to 0.19 – 0.49 s) indicated that the mean MMF amplitudes were significantly larger in the older children (*n* = 19; *M* = 0.71, *SD* = 0.45) compared to the younger children (*n* = 18; *M* = 0.40, *SD* = 0.34), *p* = .027, Cohen’s *d* = 0.77 (95% CI for Cohen’s *d*: 0.03–1.4; see Figure 3).

**Figure 2.**
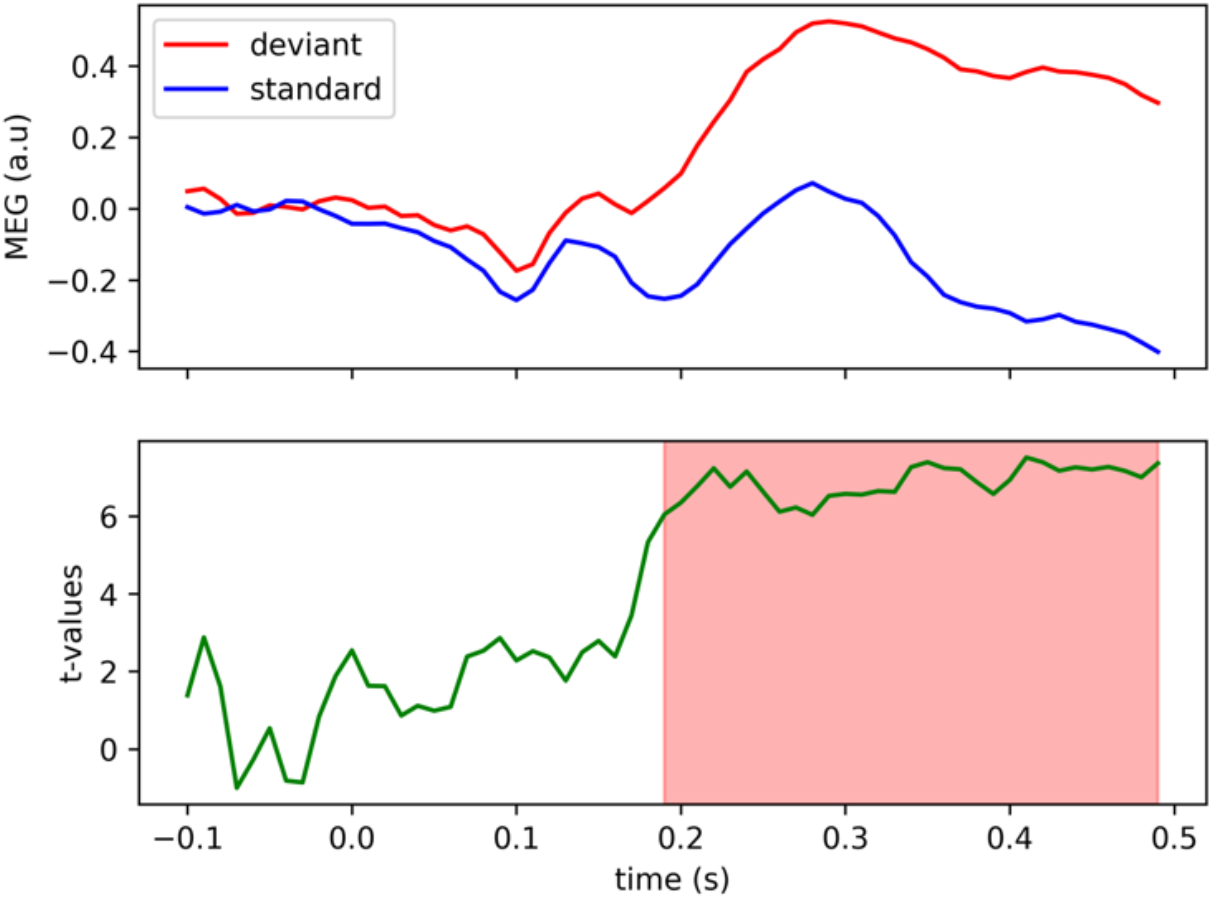
Top panel: Whole-group EMS filtered MEG data (a.u. = arbitrary units) for the standard (blue) and deviant (red) waveforms (*N* = 37). Bottom panel: results from the cluster-based permutation test. The cluster of significant t-statistics is highlighted in red.

**Figure 3.**
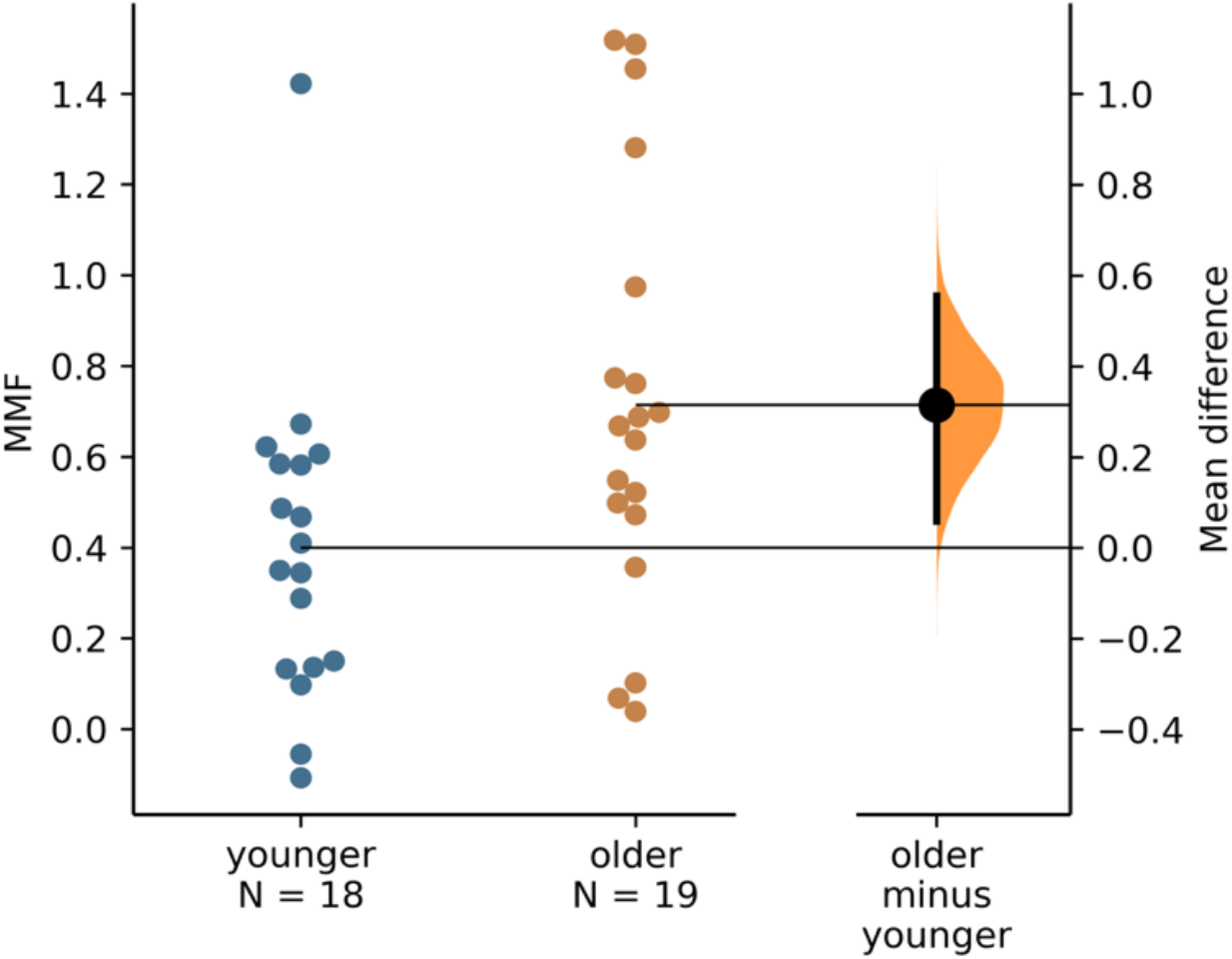
A Gardner-Altman estimation plot. Mean MMF amplitudes for all participants are plotted by age group on the two left-most axes. The effect size is presented as a bootstrap 95% confidence interval on the separate but aligned right-hand-side axis, and is displayed to the right of the data. The mean value of the older group is aligned with the effect size.

### 3.2. Source-Level Results

Across the entire sample, we found significantly larger MMF amplitudes, *p* < .05, in the three right-hemisphere ROIs: A1 (0.25–0.36 s), STG (0.25–0.36 s) and IFG (0.23–0.34 s; see Figure 4). Hence, we constrained the subsequent age-related analysis to the right-hemisphere ROIs, and to 0.23 to 0.36 s. Consistent with our hypothesis, the older children showed a significantly larger MMF amplitude, *p* < .05, in the right-IFG (0.31 to 0.33 s) relative to the younger children (see Figure 5).

**Figure 4.**
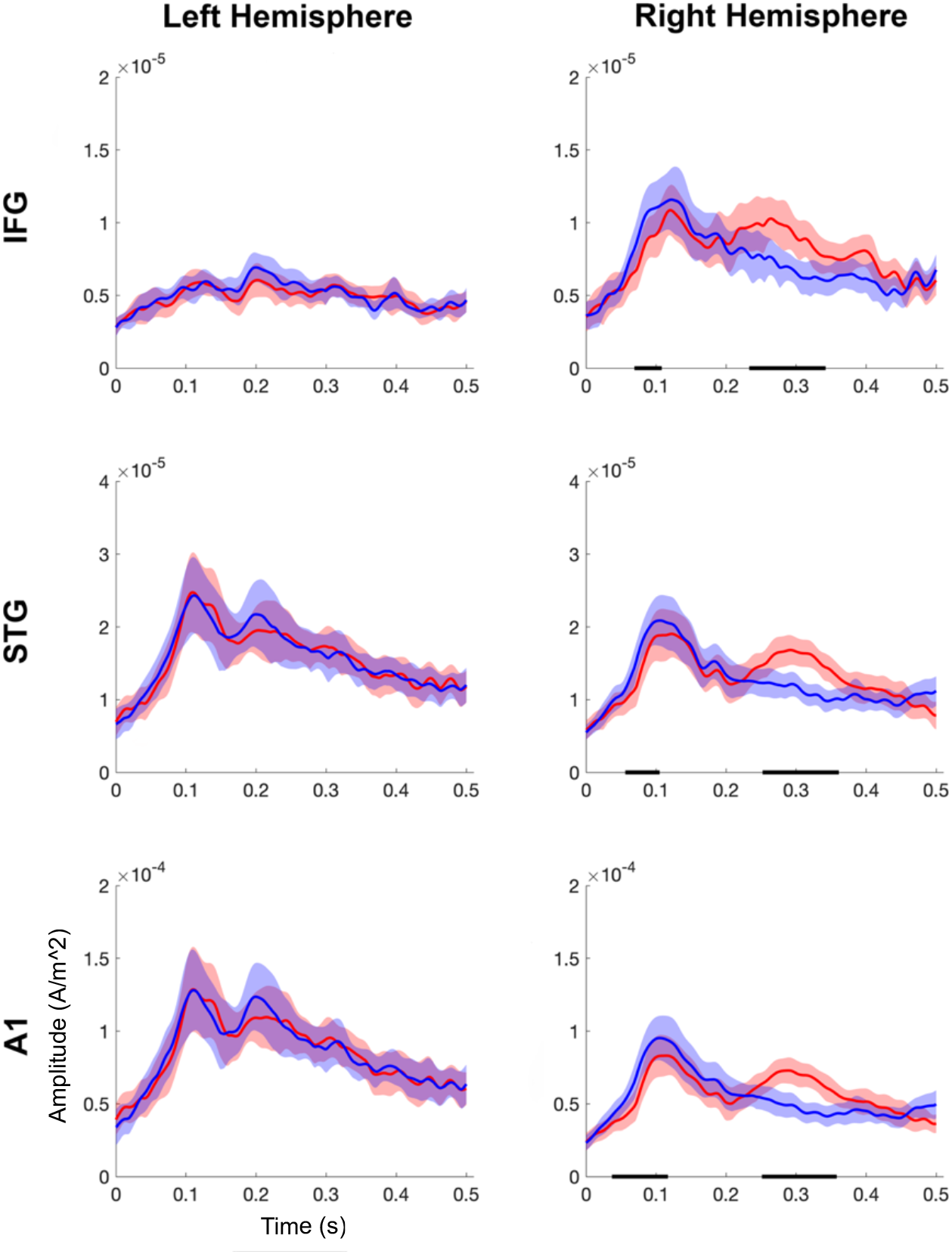
Whole-group source-level results. The two columns show the standard (blue) and deviant (red) ERF amplitudes (ampere per metre square) by time (s) for the left and right hemisphere ROIs, respectively. The shaded regions around each waveform represent 95% confidence intervals. The horizontal black line indicates clusters of significant differences (*p* < .05) between the standard and deviant ERF. Note the different y-axis scales.

**Figure 5.**
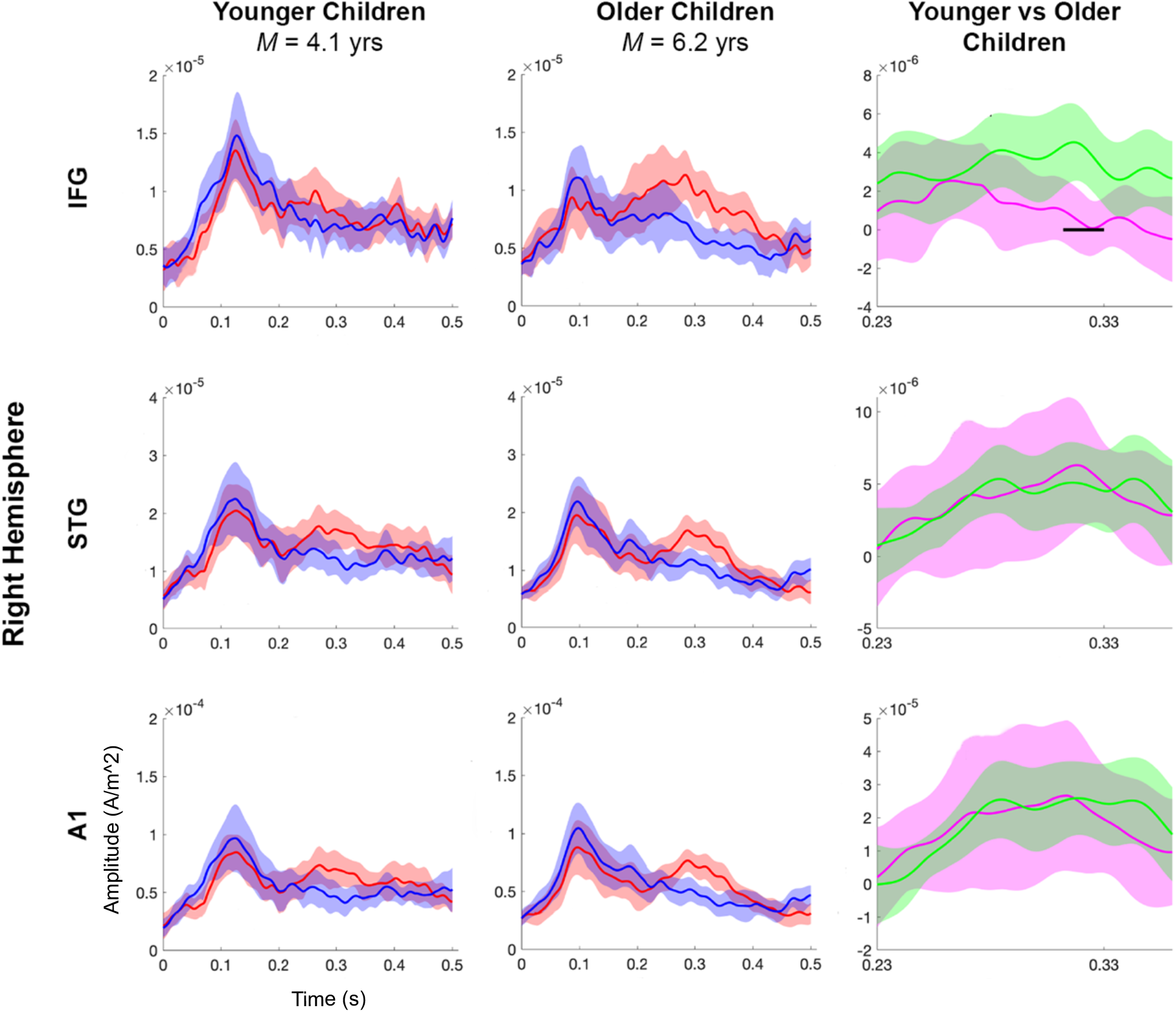
Source-level results for the younger and older children. The first two columns show the standard (blue) and deviant (red) ERF amplitudes (ampere per metre square) by time (s) for the younger and older children, respectively. The third column shows the MMF waveforms for the younger (pink) and older (green) children. The shaded regions around each waveform represent 95% confidence intervals. The horizontal black line indicates clusters of significant differences (*p* < .05) between the younger and older children’s MMF amplitudes. Note the different y-axis scales.

## 4. Discussion

This study sought to test a predictive coding account of early neurocognitive development. Consistent with our hypotheses, we found that the older children (*M* = 6.2 years, range: 5.6–6.9) showed significantly larger MMF amplitudes relative to the younger children (*M* = 4.1 years, range: 3.1–5.3). Furthermore, the older children showed a significantly larger MMF amplitude in the right inferior frontal gyrus (IFG) relative to the younger children. These findings support our proposal that predictive brain function becomes more mature across early development, and that this maturation is underpinned by increasing involvement of the frontal cortex in response to prediction errors.

### 4.1. The Maturation of the Mismatch Field

The right-lateralised findings are consistent with prior evidence that the classic, pure-tone-elicited MMN is most pronounced in the right hemisphere (Paavilainen et al., 1991). Furthermore, our finding of larger MMF amplitudes in older-relative-to-younger children is in line with the findings of previous EEG studies—both longitudinal (Chobert et al., 2014; Linnavalli et al., 2018; Putkinen, Tervaniemi, Saarikivi, de Vent, et al., 2014; Putkinen, Tervaniemi, Saarikivi, Ojala, et al., 2014) and cross-sectional (Bishop, Anderson, et al., 2011; Oades et al., 1997)—which have similarly reported an increase in the MMN amplitude across childhood. Yet our results are in conflict with the findings of several other cross-sectional EEG studies, which have reported a decrease (Csépe et al., 1992; Kraus, McGee, Carrell, et al., 1993; Kraus, McGee, Micco, et al., 1993) or no difference (Gomot et al., 2000; Kraus et al., 1999; Shafer et al., 2000, 2010) in the MMN amplitude with age.

One possible reason for the mixed findings regarding the maturation of the MMN/F may be related to variability in the methods used to calculate the MMN/F amplitude. ‘Mean measures’ (calculated by taking the average amplitude across a time window) are generally considered to be superior to ‘peak measures’ (calculated by locating the maximum amplitude within a given time window) as they are less susceptible to high-frequency noise distortions (Bishop, 2007; Bishop, Hardiman, et al., 2011; Luck, 2014b). Previous MMN studies that computed ‘mean’ measures largely reported an amplitude increase with age (Bishop, Hardiman, et al., 2011; Chobert et al., 2014; Linnavalli et al., 2018; Putkinen, Tervaniemi, Saarikivi, de Vent, et al., 2014; Putkinen, Tervaniemi, Saarikivi, Ojala, et al., 2014), whereas those using ‘peak’ measures reported either a decrease (Csépe et al., 1992; Kraus, McGee, Carrell, et al., 1993; Kraus, McGee, Micco, et al., 1993) or no difference (Shafer et al., 2000, 2010) in the MMN amplitude with age. Given the problems associated with peak measures, the mean amplitude findings seem to be the most reliable and are consistent with the findings from this study.

The key finding of this study is that older children (*M*_age_ = 6.2 years) showed significantly larger MMF amplitudes in the right IFG relative to the younger children (*M*_age_ = 4.1 years). To our knowledge, this is the first paediatric MEG study which has examined the neural sources underpinning MMN/F generation in typically developing children. It should be noted however, that—following MMN/F analysis conventions, the current source analyses were constrained to six ROIs taken from Garrido et al. (2007, 2008, 2009): the bilateral primary auditory cortices (A1), superior temporal gyri (STG), and inferior frontal gyri (IFG). We chose to constrain the analysis in this way in order to maximise the statistical power of the permutation approach (Luck, 2014a). However, there is evidence of a more extensive MMN/F neural network beyond the frontotemporal regions, including generators in the parietal cortex (Celsis et al., 1999; Molholm et al., 2005; Näätänen et al., 2019b; Opitz et al., 1999; Schall et al., 2003). As such, important ROIs may have been left out of the current source analysis. Future research should attempt to confirm the child MMN/F generators using in a considerably larger sample of participants (e.g., N =≥ 100) using functional magnetic resonance imaging (fMRI) and/or MEG whole-brain source analyses.

### 4.2. Predictive Coding Theory

We interpret the current finding of larger MMF amplitudes in older children as reflecting the maturation of predictive brain function across childhood. Specifically, we suggest that with significant physiological maturation of the brain during childhood—and in particular, the prolonged maturation of the frontal lobes (Huttenlocher & Dabholkar, 1997)— children’s brains may become increasingly proficient at extracting statistical regularities from the stream of auditory input. Equipped with this statistical knowledge, children are then able to form increasingly precise priors of the upcoming stimuli, whereby the strongest prediction would be for the presentation of a high-probability ‘standard’. The larger MMF amplitude response seen in the older children may reflect a larger prediction error response to the lower-probability ‘deviants’ relative to the precisely-predicted ‘standards’. By contrast, the relatively attenuated MMF amplitude response observed in the younger children may reflect a similar degree of prediction error elicited in response to both the less-precisely predicted ‘standards’ and ‘deviants’. Furthermore, the results support the proposal that increasingly mature predictive brain function across development is underpinned by greater involvement of the frontal cortex in responding to prediction errors.

The notion that priors become increasingly precise with age is consistent with what we intuitively understand about the differences between adult and child perception (Lucas et al., 2014). Increasingly precise prior distributions across development would progressively bias the posterior distribution (i.e., the perceptual experience) towards the narrowing prior (i.e., the brain’s stored knowledge about the world) and away from incoming sensory signals. Furthermore, increasingly precise predictions about the world have an adaptive function, resulting in fewer prediction error signals and thus, reduced demands on neural bandwidth in the long run.

Conversely, having relatively broad and less-precise priors in the early years of life would give rise to a perceptual experience that is biased towards incoming sensory signals from the world and less biased by prior knowledge. Furthermore, having imprecise, broad priors should yield more frequent prediction errors—driving more frequent model updating and thus boosting the learning rate. Indeed, it might be precisely because young children know less about the world that makes them more open to learning new information (Lucas et al., 2014).

### 4.3. Alternative Mechanistic Accounts

Predictive coding theory provides a plausible account of the current findings. While alternative mechanistic accounts of the MMN/F have been put forward, these accounts can only account for MMN/F responses elicited by auditory oddball paradigms in which the ‘standards’ are both: (a) acoustically identical to one another, and (b) repetitive in their occurrence. The ‘traditional’ auditory oddball paradigm fits these criteria, involving the presentation of a stream of repetitive standard (S) stimuli that are infrequently interrupted by the presentation of a single deviant (D) stimulus (i.e., <u>D</u>-SSS<u>D</u>-SSSSS-<u>D</u>…).

Under the ‘sensory memory’ account, a traditional MMN/F is generated when there is a detectable acoustic difference between the current input (e.g., a ‘deviant’ stimulus) and a brief memory trace of the preceding input (e.g., a ‘standard’ stimulus; Näätänen et al., 1978). Alternatively, under the ‘neuronal adaptation’ account, neurons tuned to the ‘standards’ become suppressed through repeated stimulation, whereas the rarer ‘deviant’ stimuli activate non-adapted neurons (i.e., ‘fresh afferents’), resulting in an enhanced deviant-relative-to-standard evoked response (May & Tiitinen, 2010). Finally, under predictive coding, the traditional MMN/F could reflect a relatively larger prediction error signal elicited in response to the lower-probability ‘deviants’ (P = 0.5/4) relative to the higher-probability ‘standards’ (P = 0.5). Thus, all three mechanistic accounts provide equally satisfying explanations of the traditional MMN/F.

Yet these sensory memory and neuronal adaptation accounts fail to explain MMN/Fs elicited by paradigms in which the ‘standard’ stimuli are non-repetitive in their occurrence— such as the multi-feature paradigm implemented in the current study (i.e., S-D_4_-S-D_2_-S-D_1_-S-D_4_-S-D_3…_). Under the ‘sensory memory’ account, a change-detection mechanism would signal acoustic change on every trial, resulting in equally-large evoked response amplitudes to all the stimuli and therefore, no mismatch effect. Likewise, under the neuronal adaptation account, the constant acoustic changes would not allow for habitation to any of the stimuli, again resulting in no mismatch effect.

### 4.4. Study Implications

Overall, the current neural findings offer preliminary support to the notion that predictive brain function improves across childhood. Our findings are consistent with a substantial body of behavioural evidence suggesting that young children learn the statistical regularities in their environment and use that statistical knowledge to generate predictions about future sensory events (Gopnik & Wellman, 2012; Köster et al., 2020). The predictive coding account of childhood neurocognitive development represents a relatively new and radical way of conceptualising early learning (Gopnik, 2012).

This account should also have important implications for understanding the neurocognitive underpinnings of neurodevelopmental conditions that manifest in the early childhood years. For example, predictive coding accounts of autism have proposed that both the sensory and social characteristics of autistic people may be explained by divergent precision weighting of prediction errors (Brock, 2012; Friston et al., 2013; Lawson et al., 2014, 2017; Pellicano, 2013; Pellicano & Burr, 2012; van Boxtel & Lu, 2013; Van de Cruys et al., 2013, 2014). The current findings will serve as an important baseline for testing these predictive coding accounts of autism, as well as other neurodevelopmental conditions.

### 4.5. Conclusions

In conclusion, the current study is the first to use MEG to investigate the maturation of the MMF during the childhood years. In line with our hypotheses, we found evidence of more mature predictive brain function with age, as indexed by larger MMF amplitudes in older-relative to younger children. Furthermore, this maturation appeared to be underpinned by increasing involvement of the frontal cortex in responding to prediction errors. These findings contribute to a deeper understanding of the brain function underpinning child cognitive development.

## ACKNOWLEDGEMENTS

This work was supported by the Australian Research Council Discovery Projects (DP170103148). We are extremely grateful to all the children and families who so generously gave up their time to participate in this research.

## AUTHOR CONTRIBUTIONS

**Hannah Rapaport**: Methodology, Software, Formal analysis, Investigation, Data Curation, Writing – Original Draft, Writing – Review & Editing, Visualization, **Robert A. Seymour**: Methodology, Software, Writing – Review & Editing, Visualization, **Nicholas Benikos**: Investigation, Resources, Writing – Review & Editing, **Wei He:** Methodology, Investigation, Supervision, **Elizabeth Pellicano**: Methodology, Writing – Review & Editing, Supervision, **Paul F. Sowman**: Conceptualization, Methodology, Software, Formal analysis, Writing – Review & Editing, Visualization, Supervision, Funding acquisition.

## DATA AND CODE AVAILABILITY

De-identified data and code are freely available in an Open Science Framework repository: https://osf.io/35q8n/. The MRI Estimation for MEG Sourcespace toolbox (MEMES; Seymour, 2018) can be found at https://github.com/Macquarie-MEG-Research/MEMES.

## Footnotes

1 Here the MMF was liberally defined as any significantly larger deviant-relative-to-standard amplitude within the time window of interest. Our definition stands in contrast to the definition of the classic adult MMN/F, which emerges soon after the second ‘N1’ or ‘M2’ component (Hari & Puce, 2017, p. 265; Näätänen et al., 2019a, p. 53) and is visible between 0.1 to 0.25 s following stimulus onset (Näätänen et al., 2019b). We relied on this more liberal definition as the latency of the peak mismatch effect changes across childhood (Näätänen et al., 2019a) and it would therefore have been inappropriate to constrain our analysis to the classic adult MMN time window. It should be noted that prior paediatric auditory oddball studies have likewise used liberally-defined time windows to extract MMN effects (e.g., 0.2–0.33 s in 6- to 7-year-olds, Lovio et al., 2009; 0.3–0.5 s in 4- to 12-year-olds, Partanen et al., 2013; 0.15–0.4 s in 5- to 7-year-olds, Petermann et al., 2009; 0.1–0.3 s in 9- to 13-year-olds, Putkinen et al., 2014; 0.1–0.32 s in 4- to 10-year-olds, Shafer et al., 2000).

